# OmrA sRNA Inhibits Translation of Phosphoenolpyruvate Carboxylase to Impair TCA-Cycle Flux

**DOI:** 10.64898/2026.06.26.734723

**Authors:** Thomas Stenum, Kim Boi Le Huyen, Jonas Kjellin, Sanna Koskiniemi, E. Gerhart H. Wagner, Erik Holmqvist

## Abstract

Small RNAs (sRNAs) rarely cause strong growth phenotypes upon overexpression, complicating efforts to link regulatory interactions to physiological outcomes. Here, we report that high levels of the *Escherichia coli* sRNA OmrA, but not its sibling OmrB, severely inhibit growth in glucose minimal medium. Genetic, biochemical, and physiological analyses indicate that OmrA-dependent toxicity results from reduced flux through the tricarboxylic acid (TCA) cycle. A UV-based suppressor screen identified mutations in the gene encoding Hfq, the RNA-chaperone that aids sRNA-mRNA interactions. Secondly, three independent mutations clustered in the ribosome-binding site of *ppc*, encoding phosphoenolpyruvate carboxylase, a key anaplerotic enzyme. OmrA directly inhibits Ppc translation via Hfq-dependent base-pairing in the *ppc* 5′ UTR, including the mutated nucleotides obtained in the genetic screen. OmrA is significantly more effective than OmrB in *ppc* repression *in vivo* and *in vitro*, consistent with sequence divergence in their central regions. Supplementation with glutamate, glutamine, or downstream TCA cycle metabolites fully restores growth, linking reduced Ppc levels to metabolic limitation. These results identify *ppc* as a physiologically relevant OmrA target and suggest how RNA toxicity can uncover central metabolic nodes used by sRNAs to modulate bacterial physiology.

## Introduction

RNA toxicity is a phenomenon observed in both bacterial and eukaryotic systems. It generally implies that either the overexpression of a certain RNA, or the presence of a mutated version of an endogenous RNA, can cause harmful effects such as growth arrest or inviability. In humans, many severe pathologies are the result of mutations, often involving repeat expansions, that create aberrant RNAs, which via altered protein binding or localization cause misregulation. For instance, in myotonic dystrophy, CUG or CCUG expansions in noncoding mRNA regions cause sequestration of splicing factors, and ultimately splicing defects (Sicot and Gomes-Pereira, 2013). In another class of disease-causing mutations, found in hereditary neurodegenerative disorders, long CAG trinucleotide sequence expansions in an mRNA generate so-called polyQ/glutamine proteins (Tandon *et al*., 2024). The severity of disease is caused by toxic inclusion bodies that affect many key pathways. Toxicity is thus here only indirectly caused by the aberrant mRNA, and disease is ultimately caused by the faulty proteins.

In bacteria, observations in several labs have shown that overexpression of some RNAs can compromise growth rate or even viability under certain growth regimens, irrespective of whether an RNA is translatable/ translated or not (see below). One study demonstrated that some mutant versions of the *gfp* gene conferred toxicity in *Escherichia coli* (*E. coli*), and that this effect was caused by the mutant RNAs produced and entirely independent of GFP translation (Mittal *et al*., 2018). The results in this paper failed to identify a mechanism that could explain the strong phenotypes observed.

A major class of regulatory RNAs in bacteria and archaea, sRNAs, has been intensively studied for many years (Holmqvist and Wagner, 2017; Hör *et al*., 2020). Most sRNAs fail to elicit strong effects on growth upon deletion or overexpression, making it challenging to use phenotypes to guide mechanistic/ functional understanding. This does not necessarily imply that most sRNAs are dispensable for bacterial survival, but rather that conditions in which a particular sRNA has a strong impact on bacterial proliferation are difficult to predict. Nevertheless, in a few cases, sRNA overexpression entails strong growth effects. For instance, the DNA damage-induced sRNA OxyS, when highly expressed from plasmids, is toxic, apparently via repression of NusG translation. Lower NusG levels result in readthrough of Rac prophage genes, causing expression of the KilB protein which - via effects on FtsZ - causes growth arrest to enable subsequent DNA repair (Barshishat *et al*., 2018). A second example comes from a study of the sRNA Spot 42. As part of the catabolite repression response, this sRNA downregulates mRNAs of genes relevant to the use of non-glucose carbon sources. Overexpression of Spot 42 significantly decreased the cell growth on non-preferred carbon sources (Beisel and Storz, 2011). In *Mycobacterium tuberculosis*, overexpression of the sRNA 6C is lethal, and growth in *M. smegmatis* is decreased (Arnvig and Young, 2009; Mai *et al*., 2019). In *E. coli*, the Qin phage-encoded DicF sRNA, a regulator of *ftsZ*, renders cells unable to grow on xylose-only medium, due to repression of *xylR* (Balasubramanian *et al*., 2016). Thus, under certain conditions, sRNAs can confer toxicity.

Many bacteria encode so-called sibling sRNAs (Caswell *et al*., 2014), presumed to originate from gene duplication events. We have previously investigated two paralogous *E. coli* sRNAs, OmrA and OmrB, with respect to their multiple antisense targets and their impact on bacterial physiology, in particular in decisions to enter/ maintain a motility or biofilm program (Holmqvist *et al*., 2010; Wagner and Romby, 2015; Hoekzema *et al*., 2019; Romilly *et al*., 2020). Other labs had earlier shown that these RNAs repress many mRNAs encoding outer membrane proteins (Guillier and Gottesman, 2006; Guillier and Gottesman, 2008). Almost all known targets are regulated by both OmrA and OmrB; their almost identical (one nt mismatch) 5’ tails appear to account for the observed effects. By contrast, the central regions in these two RNAs have over evolutionary time acquired major changes, suggesting them to be on a trajectory to acquire distinct target profiles. Additionally, even though both genes are regulated by the transcription factor OmpR (Guillier and Gottesman, 2006; Brosse *et al*., 2016), only OmrA is specifically transcribed by Sigma S-containing RNA polymerase (Peano *et al*., 2015). So far, targets unique to only one of these sRNAs appear to be rare, but a recent paper demonstrated that OmrA, but much less so OmrB, represses translation of the *btuB* mRNA that encodes an importer of cobalamine (Bastet *et al*., 2024).

In experiments aimed at finding targets that responded differently to overexpression of either OmrA or OmrB, we inadvertently found that while growth was unaffected in rich medium, high levels of OmrA, but not OmrB, severely impeded growth in glucose-minimal media (see below). This suggested that OmrA was (conditionally) toxic upon overexpression. Given that OmrA is a noncoding RNA, we envisioned mechanisms that in principle might apply: 1) one in which high levels of an sRNA by antisense may downregulate target mRNAs whose gene products are needed in the physiological conditions tested, 2) one in which an sRNA titrates an RNA-binding protein which is conditionally essential for growth, or 3) though less likely, even binding of an sRNA to a second sRNA (“spongeing”; e.g. (Figueroa-Bossi and Bossi, 2018)) could indirectly impact various processes.

In this paper, we address the mechanism of OmrA toxicity and identify the pathway affected. Our results suggest that the difference between the middle regions of the sibling RNAs tentatively accounts for OmrA toxicity. We furthermore show that the observed growth defect can be traced to central carbon metabolism. OmrA targets *ppc* mRNA, encoding phosphoenolpyruvate carboxylase, an enzyme that replenishes oxaloacetate to maintain the flux of intermediates through the TCA cycle.

## Materials and methods

### Oligodeoxyribonucleotides

All oligodeoxyribonucleotides used in the study were purchased from Eurofins Genomics and are listed in Supplementary Table S1.

### Bacterial strains, media and growth conditions

Strains used in this study are listed in Supplementary Table S2. *E. coli* K-12 strain MG1655 was used for all experiments except for Northern blot analysis, where *E. coli* MC4100 was used. The strain TOP10 (Invitrogen) was used for construction and propagation of plasmids. Cultures were grown at 37°C shaking at 180 rpm in M9 media supplemented with 0.2% glucose and 10 μg/ml thiamine, or 0.1% casamino acids (CAA), unless otherwise specified. Individual amino acids were added at 50 μg/ml. Antibiotics were added at the following concentrations: ampicillin 100 μg/ml, chloramphenicol 30 μg/ml, kanamycin 50 μg/ml.

### Strain constructions

Chromosomal manipulations were done by λ-red recombineering using the pSIM5 plasmid series (Datta *et al*., 2006). The *omrAB* locus was replaced by a PCR-generated fragment (primers EHO-245/EHO-246) containing a tetracycline resistance gene flanked by FRT sites. This construct was further moved into a fresh wt background by P1 transduction, and confirmed by PCR (EHO-247/ EHO-249) and sequencing. Finally, the tetracycline cassette was removed by FLP-mediated recombination using plasmid pCP20 (Cherepanov and Wackernagel, 1995). C-terminal 3xFLAG-tagging of *ppc* was done using a PCR fragment (EHO-2307/ EHO-2308) amplified from pSUB11 (Uzzau *et al*., 2001). The *ppc*-3xFLAG construct was moved into a fresh wt background by P1 transduction, validated by PCR (EHO-2309/ EHO-2310) and sequencing.

### Plasmid constructions

Plasmids used are listed in Supplementary Table S3. pKB009 was constructed through PCR-based site-directed mutagenesis of pEH67 (Holmqvist *et al*., 2010) using two partially overlapping primers (EHO-2370/EHO-2371). The resulting PCR product was treated with *Dnp*I for 1 h at 37°C before transformation into TOP10 cells. Successful mutagenesis was confirmed by PCR (EHO-1006/ EHO-1122) and sequencing. pTSS58 (*ppc::gfp*) was constructed by inserting a *Nsi*I/*Nhe*I-digested PCR product (EHO-2287/EHO-2288) into *Nsi*I/*Nhe*I-digested plasmid pXG-10 (Urban and Vogel, 2007). Plasmids pTSS59 (*ppcM1::gfp*), pTSS60 (*ppcM2::gfp*), pTSS61 (*ppcM3::gfp*), and pKB013 (*ppcM1*::gfp*) were created by site-directed mutagenesis using primers EHO-2289/EHO-2290, EHO-2291/EHO-2292, EHO-2293/EHO-2294, and EHO-2470/EHO-2471, respectively, using pTSS58 as template. Successful mutagenesis was confirmed by PCR (EHO-827/ EHO-828) and sequencing.

### UV-mutagenesis and whole genome sequencing

Ten individual cultures of MG1655 harboring plasmid pEH67 were grown o.n. in M9 medium supplemented with 0.2% glucose, 0.1% CAA, and 100 μg/ml ampicillin. Overnight cultures were pelleted, resuspended in M9 medium supplemented with 0.2% glucose, 10 μg/ml thiamine, and 100 μg/ml ampicillin, and incubated at 37 °C with shaking for 30 min. Two ml of each culture was spotted on a plastic tray and treated with UV-light (10 mJ/cm^2^), transferred to 10 ml glass tubes, incubated in the dark for 10 min at 37 °C, and spread on M9 agar plates supplemented with 0.2% glucose, 10 μg/ml thiamine, and 100 μg/ml ampicillin. After 30-40 hr incubation at 37 °C, eight large colonies per plate were re-streaked twice on identical plates. To discard suppressors linked to mutations in the plasmid-encoded *omrA* gene, the pEH67 plasmid was extracted from each fast-growing clone, retransformed into the wild-type strain, and assayed for toxicity on M9 agar plates supplemented with 0.2% glucose, 10 μg/ml thiamine and 100 μg/ml ampicillin. Suppressor strains associated with non-toxic pEH67-derivatives were discarded. High molecular weight gDNA was prepared from the remaining suppressor strains using the Wizard HMW DNA Extraction kit (Promega). The DNA was converted to barcoded sequencing libraries using the Rapid Barcoding Kit 96 V14 (SQK-RBK114.96) which were sequenced on an Oxford Nanopore MinION Mk1C instrument using an R10.4.1 (FLO-MIN114) flowcell. Nanopore duplex reads were aligned to the reference genome NC_000913.3 using minimap2 (v2.24) and SAM files were converted to sorted and indexed BAM files using samtools (v1.17). Variant calling was performed to produce a compressed and indexed VCF file, which was subsequently filtered prior to downstream analyses. Variant data (chromosome, position, reference and alternate alleles, depth of coverage, and allele frequency) were extracted using bcftools query and formatted into BED-like tables. Gene coordinates were derived from a GFF annotation file using gawk, which produced a BED file containing chromosomal positions and corresponding gene names. Variants were intersected with these genomic features via a custom AWK script that identified variants located within gene annotations or in intergenic regions. For each position, the nearest upstream and downstream genes were also reported.

### Northern blot

Total RNA was purified using the hot phenol method as described in (Holmqvist *et al*., 2018). RNA samples were mixed 1:1 in GLII loading buffer (0.025 % w/v xylene cyanol, 0.025 % w/v bromophenol blue in formamide), denatured at 95°C, and separated on polyacrylamide (PAA)/ 8 M urea gels along with radioactively labelled size marker pUC19 MSP1 (ThermoFischer). RNA was transferred to a nitrocellulose Hybond-N+ membrane (Amersham, Cytiva) by wet electroblotting at 4°C for 2 h at 360 mA, crosslinked at 1200 mJ UV light, and prehybridized in Church buffer (0.25 M phosphate buffer pH 7.2, 1 mM ethylenediaminetetraacetic acid solution (EDTA), 7% sodium dodecyl sulfate (SDS)) (Church and Gilbert, 1984) for 45 min at 42°C. A 5’-^32^P-labeled DNA oligonucleotide was hybridized to the membrane for 2-24 h. Membranes were washed three times in 0.5 × saline-sodium citrate (SSC)/0.1% SDS. Dried membranes were exposed to a phosphor screen and radioactive signals detected with a Typhoon phosphorimager (Cytiva).

### GFP fluorescence measurements

Bacterial cells grown o.n. in LB medium at 37°C were diluted 1:100 in fresh M9 medium supplemented with 0.2% glycerol and 0.1% CAA. GFP measurements were performed in 96-well microtiter plates in a TECAN Infinite Pro plate reader as described (Rizvanovic *et al*., 2021).

### Toxicity plating assays

Bacterial cells grown o.n. in LB medium at 37°C were washed and diluted in PBS to OD600 = 0.3, and serially diluted tenfold. Five μl of each dilution was spotted on LB agar plates or M9 plates supplemented with 0.2% glucose and 10 μg/ml thiamine, or 0.1% CAA, and incubated o.n. at 37°C.

### Hfq protein purification

Hfq was purified from *E. coli* strain BL21(DE3)pLysS harboring the Hfq-6xhis-encoding plasmid pTE607 (Fender *et al*., 2010). Cells were grown o.n. in LB containing 50 μg/ml ampicillin and 30 μg/ml chloramphenicol at 37°C, diluted 1:50 in 500 ml of fresh medium, grown until OD600 = 0.7, after which IPTG was added to 0.5 mM and growth continued for 3 h at 30°C before harvest by centrifugation.

Pellets were resuspended in 2 ml of lysis buffer (20 mM Tris-HCl pH 7.5, 0.5 M NaCl, 10% (v/v) glycerol, 0.1% (v/v) Triton X-100, 100 U DNase I, 10 mM imidazole) and lysed in a BeadBugTM-6 microtube homogenizer (Benchmark Scientific) at 4.0 m/s for 20 sec, followed by 15 min centrifugation at 13,000 g in a cold room. The supernatant was heated at 80°C for 15 min, insoluble material removed by centrifugation, and filtered through a 0.45 μm MillexHA filter (Millipore). The lysate containing 6xhis-tagged Hfq was mixed with 1:5 of volume of 50% (v/v) Ni-NTA Magnetic Agarose beads (Qiagen), equilibrated in 20 mM Tris-HCl pH 7.5, 0.5 M NaCl, 10% (v/v) glycerol, 10 mM imidazole, and shaken for 1 h at 4°C. Beads were washed twice with 500 µl of wash buffer (20 mM Tris-HCl pH 7.5, 0.7 M NaCl, 1 M imidazole) using magnetic separator between washes. The beads were resuspended in 100 µl of elution buffer (20 mM Tris-HCl pH 7.5, 0.3 M NaCl, 0,5 M imidazole), incubated for 1 min before collecting the eluate. The eluted sample was dialyzed o.n. in storage buffer (50 mM Tris-HCl pH 7.5, 1 mM EDTA, 5% glycerol, 50 mM NH_4_Cl, 0.1% (v/v) Triton X-100) using Slide-A-Lyzer™ Dialysis Cassettes, 7K MWCO (Thermo Scientific) and stored at 4°C. Concentrations were determined by the Bradford method (Bio-rad) with BSA as standard. Hfq molar concentrations were calculated on the basis of hexamers.

### In vitro transcription and labelling of RNA

OmrA and OmrB RNAs and mutant versions thereof were generated as previously described (Holmqvist *et al*., 2010). To generate *ppc* mRNA fragments (from -88 to +40, or -88 to +87), DNA templates containing a T7 promoter sequence were generated by PCR using primers sets EHO-2312/EHO-2313 and EHO-2312/EHO-2470 for EMSA and toeprint assays, respectively, with *E. coli* MG1655 DNA as template. The PCR products were used for *in vitro* transcription following the MEGAscript kit’s instruction (Life Technologies). RNAs were purified and 5’ end-labelled as described (Eleftheraki and Holmqvist, 2024). Labelled RNAs were purified using MicroSpinTM G-50 columns (Cytiva, Amersham).

### Electromobility Shift Assays

*In vitro* transcribed RNA dissolved in sterile water was denatured at 95°C for 1 min, cooled on ice for 2 min, diluted in binding buffer (25 mM Tris-HCl pH 7.4, 100 mM NaCl, 1 mM MgCl2), and renatured at 37°C for 5 min. 5’-labelled *ppc* mRNA (1 nM final concentration) was mixed with an increasing concentration of sRNA, and incubated at 37°C for 30 min. Samples were loaded on native 6% polyacrylamide gels in 0.5 x TBE buffer, and eletrophoresed at 200 V in a cold room. Radioactive signals were detected using a Typhoon PhosphorImager (Cytiva).

### Toeprint assays

Toeprint assays were carried out as in (Romilly *et al*., 2020). Final concentrations of reaction components were: 20 nM *ppc* mRNA, 100 nM 30S subunits, 300 nM initiator tRNA, 0.5 mM dNTPs, 500 nM Hfq, and 10 µM sRNAs (OmrA or ArcZ). Sequence ladders were produced according to the USB Thermo Sequenase Cycle Sequence kit’s instruction (Affymetrix), with a *ppc* PCR product (EHO-2312/ EHO-2470) as template.

### Western blotting

Bacterial o.n. cultures were diluted 1:50 in fresh M9 medium with 0.1% CAA and grown at 37°C shaking at 180 rpm for 3 h (exponential phase). Cells were pelleted by centrifugation, resuspended in Laemmli Sample Buffer, denatured for 5 min at 95°C, separated on Mini-PROTEAN TGX Stain-Free protein gels (BioRad), and transferred to PVDF membranes using the TransBlot TURBO transfer system (BioRad). Membranes were blocked with 5% milk powder in TBS-T for 1 h at r.t.. FLAG-tagged Ppc was detected using an HRP-conjugated anti-FLAG antibody at 1:10,000 dilution (Sigma), and GroEL with an HRP-conjugated anti-GroEL antibody (Sigma) at 1:50,000 dilution. Chemiluminescence was detected using Amersham ECL Prime reagents according to the supplier’s protocol (Cytiva) on a ChemiDoc™ System (BioRad).

## Results

### OmrA sRNA overexpression causes growth inhibition

Plasmid overexpression is a common approach to amplify the effects of sRNAs when characterizing their functions. In most cases, sRNA overexpression fails to cause obvious phenotypes during growth in standard laboratory media. E.g., overexpression of sRNAs OmrA and OmrB does not affect growth of *E. coli* on LB agar plates, compared to a strain harboring an empty vector (Fig. 1A). By contrast, when grown on minimal M9 medium agar plates supplemented with glucose (M9-glu), an OmrA overexpression strain gave smaller colonies (Fig 1A). Despite the high sequence similarity between OmrA and OmrB, particularly their almost identical “seed” regions (Guillier and Gottesman, 2008), overexpression of OmrB barely reduced growth on M9-glu. To quantify the OmrA-dependent growth defect, strains were grown in LB, washed and serially diluted in PBS, and spotted on agar plates with LB medium, M9-glu, or M9 medium supplemented with casamino acids (M9-CAA). Overexpression of OmrA did not affect growth on LB plates, and only slightly reduced CFUs on M9-CAA (Fig. 1B). On M9-glu plates, OmrA overexpression resulted in a ten-thousand-fold reduction in CFUs, compared to the empty vector control (Fig 1B). By contrast, overexpression of OmrB did not affect CFUs on any of the media, but resulted in a small colony phenotype on M9-glu plates, indicative of a slight reduction in growth rate (Fig. 1B). Hence, overexpression of OmrA is toxic under conditions when glucose serves as the sole carbon source.

**Figure 1.**
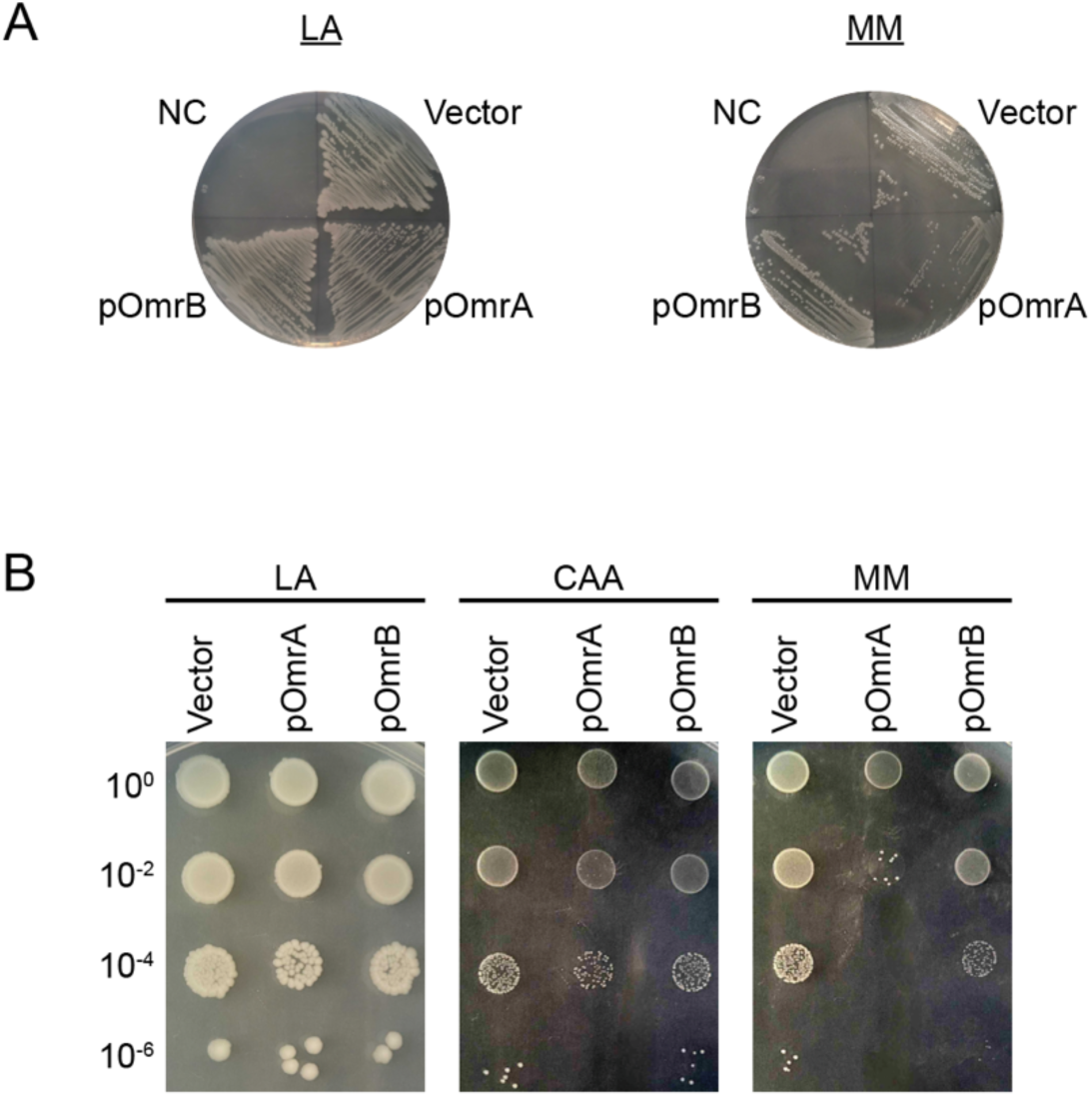
Overexpression of OmrA, but not OmrB, causes a slow-growth phenotype in minimal growth medium. (A) *E. coli* MG1655 Δ*omrAB* transformed with plasmids constitutively overexpressing OmrA (pOmrA), OmrB (pOmrB), or an empty control plasmid (Vector) were streaked on Luria-agar plates (LA, left) or M9 miminal medium supplemented with glucose (MM, right). NC: negative control where no bacteria was streaked. (B) The same strains as described in A were grown overnight in LB medium, subjected to tenfold dilutions, and spotted on plates with Luria-agar (LA, left) or M9 miminal medium supplemented with glucose and casamino acids (CAA, middle), or M9 miminal medium supplemented with glucose (MM, right).

### OmrA-dependent toxicity is coupled to the TCA cycle

In agreement with the results from growth on solid media, overexpression of OmrA strongly reduced growth in liquid M9-glu medium (Fig. 2A). Again, addition of casamino acids completely abolished this phenotype (Fig. 2A), suggesting that OmrA-dependent toxicity may be coupled to limitation of one or several amino acids. Congruent with this, a mix of all twenty amino acids phenocopied the effect of casamino acids (Fig. 2A). The amino acids were divided into two bins of ten each (mix A and B). While mix A somewhat increased the growth rate of the OmrA-overexpression strain, mix B fully restored growth to that of the empty vector (Fig. 2A). Next, the effect of OmrA overexpression was monitored with each of the amino acids in mix B added separately. Strikingly, addition of glutamate or glutamine fully restored the growth rate of the OmrA overexpression strain (Fig. 2B). These two amino acids are synthesized from the TCA-cycle metabolite α-ketoglutarate, indicating that OmrA-dependent toxicity is linked to the metabolic flow through the TCA cycle.

**Figure 2.**
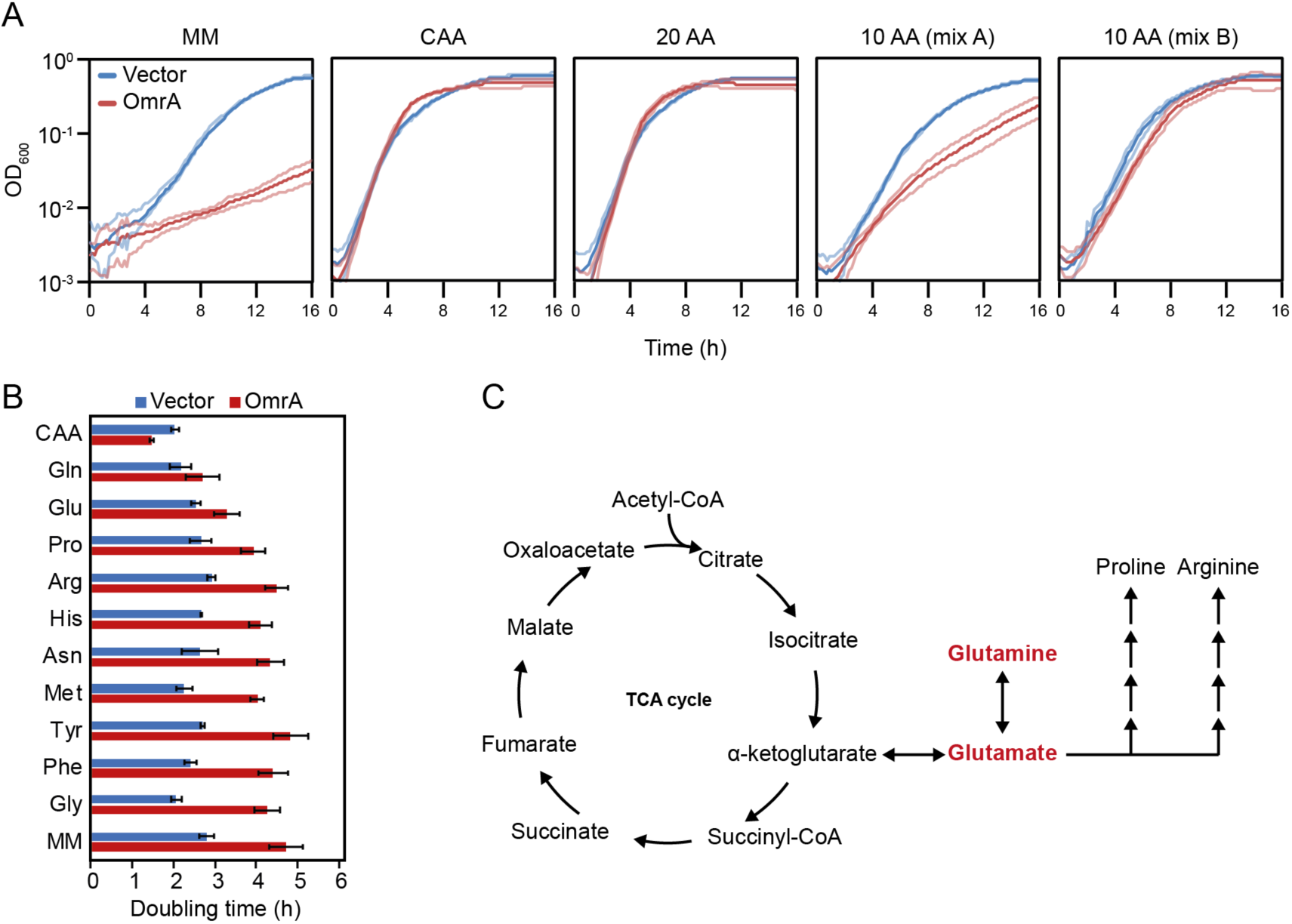
OmrA-dependent growth deficiency is linked to amino acid starvation. (A) *E. coli* MG1655 Δ*omrAB* harboring an OmrA overexpression plasmid, or its empty counterpart, were grown overnight in LB medium and diluted 100-fold in M9 minimal medium supplemented with glucose (MM), glucose and casamino acids (CAA), glucose and all twenty amino acids (20 AA), glucose and ten of the amino acids (10 AA (mix A)), or glucose and the remaining ten amino acids (10 AA (mix B)). See Materials and methods for specification of amino acid composition in mix A and B, respectively. Growth of the diluted cultures was monitored by measuring optical density (OD_600_) for sixteen hours in a 96-well plate reader. Bright blue or red lines show the mean growth rate and faint lines show standard deviation based on four biological replicates. (B) The same strains as described in A were grown overnight in LB medium and diluted 100-fold in MM supplemented with casamino acids, or a single amino acid from mix A, as indicated in the figure. Growth was monitored in 96-well plates as described in A during sixteen hours. For each strain and condition, the maximum growth rate was calculated (Materials and methods). Bars show the mean growth rate, and error bars the standard deviation based on four biological replicates. (C) Schematic showing the main metabolic intermediates of the TCA cycle, including the pathway from α-ketoglutarate to glutamate and glutamine.

### OmrA-dependent toxicity is due to decreased flux through the TCA cycle

Since glutamate and glutamine appeared to be limiting upon OmrA overexpression, we wondered whether OmrA reduces the expression of an enzyme in the glutamate/glutamine biosynthetic pathway. To this end, the OmrA overexpression strain was transformed with plasmids expressing each enzyme on the route from acetyl-coenzyme A to glutamate: GltA, AcnB, IcdA, GdhA, and AlaA (Fig. 3A). However, none of these enzymes restored the growth phenotype (Fig. 3B). By contrast, addition of the TCA cycle metabolites α-ketoglutarate, malate, or fumarate, restored growth. Pyruvate, which is at the entry point of the TCA cycle, did not (Fig. 3C). Together, this indicated that OmrA-dependent growth inhibition depends on glutamate/glutamine limitation due to decreased flow through the TCA cycle. Since overexpression of TCA cycle enzymes could not restore growth (Fig. 3B), the reduced flow through the TCA cycle may rely on halted replenishment of TCA cycle metabolites.

**Figure 3.**
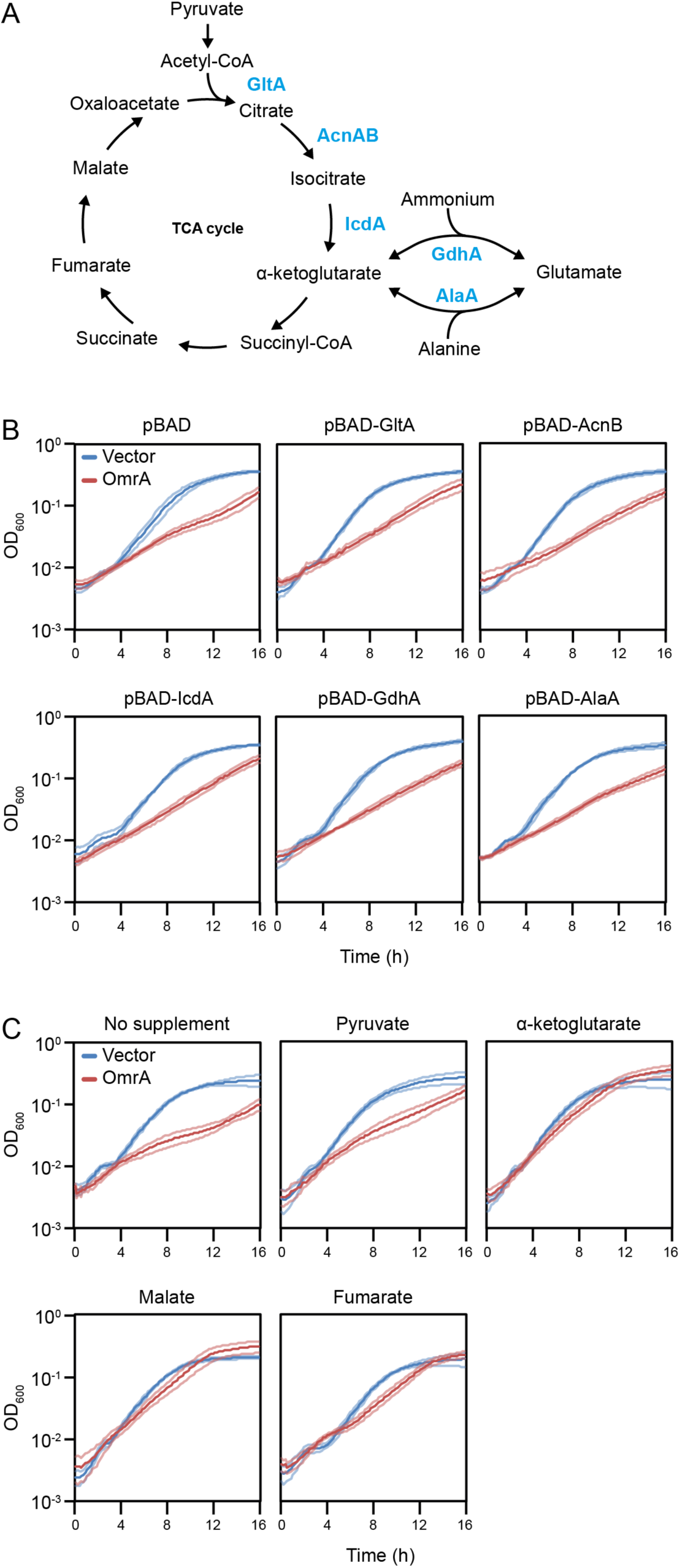
OmrA-dependent growth deficiency is linked to reduced metabolic flow in the TCA cycle. (A) Schematic showing the TCA cycle and the required enzymes to convert pyruvate to glutamate. (B, C) *E. coli* MG1655 Δ*omrAB* harboring an OmrA overexpression plasmid, or its empty counterpart, were grown overnight in LB medium and diluted 100-fold in M9 minimal medium supplemented with glucose. Growth was followed in a 96-well plate reader for sixteen hours by measuring optical density at 600 nm (OD_600_). In (B) each strain harbored an additional plasmid conferring overexpression of the indicated TCA cycle enzymes. In (C) the media were supplemented with the indicated TCA cycle metabolites. Bright blue or red lines: mean growth rate. Faint lines: standard deviation based on four biological replicates.

### UV-based mutagenesis identifies suppressors of OmrA toxicity

In an attempt to identify the direct cause of OmrA-dependent toxicity, suppressors of the growth phenotype were selected as fast-growing colonies on M9-glu plates after UV irradiation (Fig. 4A). To discard trivial suppressors due to mutations in the *omrA* gene, the OmrA-overexpression plasmids were extracted from each suppressor strain, re-transformed into the wild-type strain, and scored for growth on M9-glu plates. Only suppressor strains that harbored plasmids retaining the ability to cause the toxic phenotype were kept for further analysis. To identify suppressor mutations on the chromosome, 23 suppressor strains were subjected to whole genome sequencing. In total, almost 200 mutations were identified, with 22 strains having mutations in more than one gene (Supplementary Table S4). Eleven of the suppressor strains had mutations in the *hfq* gene, encoding the Hfq protein that mediates interactions between sRNAs and their targets (Vogel and Luisi, 2011). The mutations spanned all three major RNA-binding faces of Hfq; proximal, distal face, and rim (Fig. 4B). To understand the role of Hfq in OmrA-dependent toxicity, we first monitored the effect of Hfq on OmrA expression. To this end, OmrA levels were monitored by Northern blot in strains expressing previously characterized Hfq mutants (Mikulecky *et al*., 2004; Sauer and Weichenrieder, 2011; Sauer *et al*., 2012; Zhang *et al*., 2013), carrying point mutations in each of the three RNA-binding faces. Compared to the wild-type strain, a mutation in the proximal face of Hfq (F42A) strongly reduced OmrA levels (Fig. 4C). In agreement with this, the OmrA-dependent toxicity was abolished when Hfq carried the F42A mutation, or upon *hfq* gene deletion (Fig. 4D). In contrast to F42A, the distal face mutation Y25D resulted in increased OmrA levels (Fig. 4C). Despite this, the Y25D mutation suppressed OmrA-dependent toxicity (Fig. 4D). Since the distal face typically binds mRNA targets to mediate sRNA-mRNA binding, this suggests the toxicity to be the result of OmrA-dependent regulation of an mRNA target. The rim mutation R16A did not affect OmrA levels and failed to suppress toxicity. Together, these results show that Hfq is required for OmrA-dependent toxicity, either by stabilizing OmrA through the proximal face, or by a putative interaction with an mRNA regulated by OmrA.

**Figure 4.**
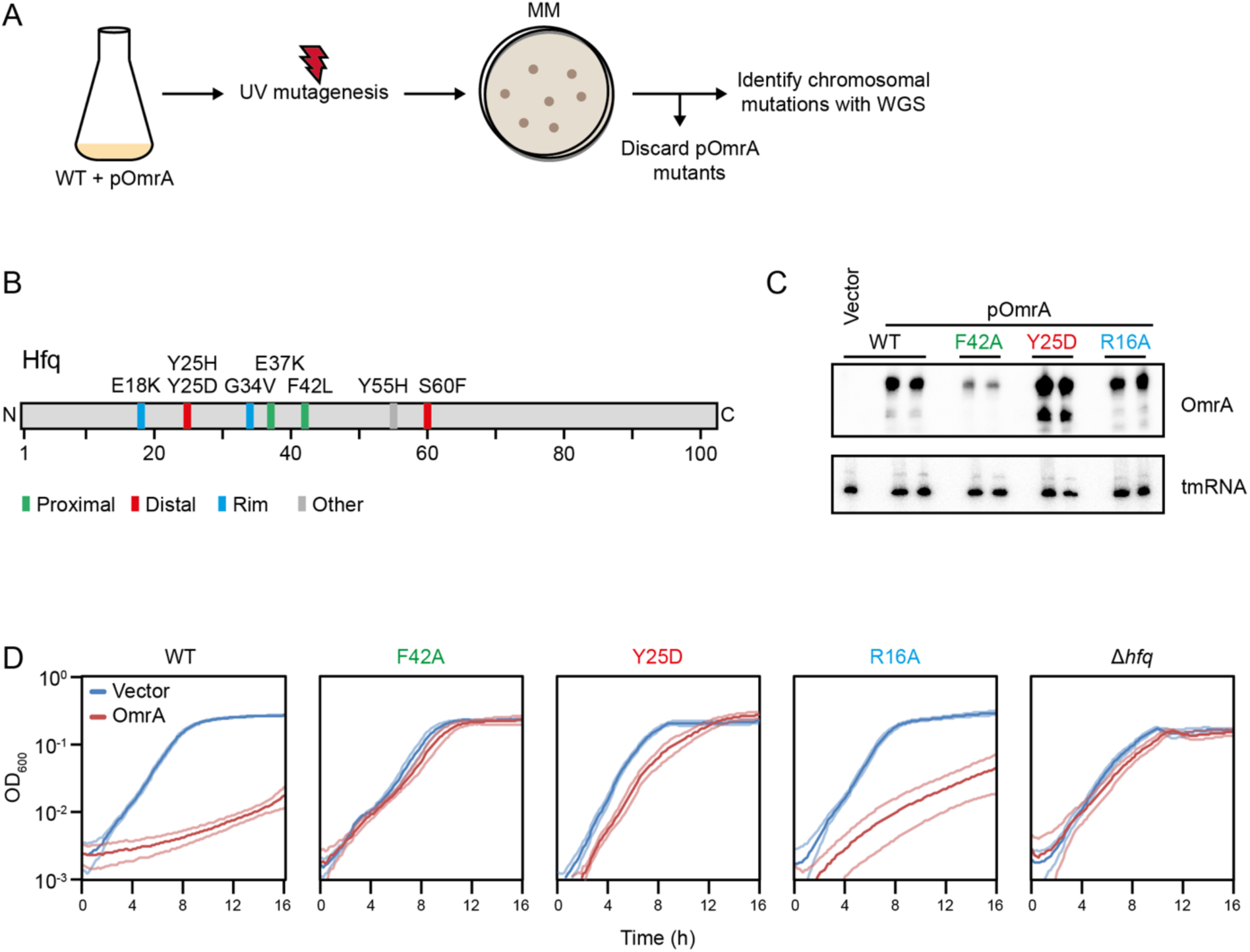
Mutations in Hfq suppress OmrA-dependent growth deficiency. (A) Schematic showing the mutagenesis strategy used to identify chromosomal mutations that suppress OmrA-dependent growth deficiency. For a detailed description, see Materials and methods. (B) Non-synonymous suppressor mutations identified in the Hfq protein. Identified amino acid substitutions and their positions in the major RNA-binding domains of Hfq are shown. (C) Northern blot analysis of OmrA expression levels in strains with previously characterized mutations in the three major RNA-binding sites of Hfq. Probing for 5S ribosomal RNA was used as loading control. (D) The wild-type strain and the indicated isogenic *hfq* mutants were grown in M9-glu medium for sixteen hours in a 96-well plate reader. Each strain harbored the OmrA overexpression plasmid or an empty control plasmid. Growth was monitored by measuring OD_600_. Bright blue or red lines: mean growth rate. Faint lines: standard deviation based on four biological replicates.

### OmrA inhibits synthesis of the metabolic enzyme Ppc

Beyond *hfq,* three suppressor strains had clustered independent mutations in *ppc*, encoding the metabolic enzyme phosphoenolpyruvate carboxylase (Ppc). Ppc feeds the TCA cycle by directly converting phosphoenolpyruvate to oxaloacetate, thereby by-passing pyruvate and acetyl-coenzyme A (Fig. 5A) (Peng *et al*., 2004). In the absence of Ppc, *E. coli* cells are unable to grow on M9-glu (Coomes *et al*., 1985; Tong *et al*., 2020). Identification of suppressor mutations in *ppc* is consistent with the lack of growth resumption upon addition of pyruvate (Fig. 3C), which mainly aids in synthesis of acetyl-CoA rather than in downstream effects on oxaloacetate. The suppressor library included three independent point mutations in the *ppc* gene, located ten, eight, and five nucleotides upstream of the *ppc* start codon, respectively (Fig. 5B). To investigate whether the mutations affected Ppc synthesis, we used a translational fusion comprising the *ppc* 5’UTR and the first 20 codons of the *ppc* ORF, cloned in-frame with *gfp*. The gene fusion was expressed from a heterologous promoter on a plasmid. Compared to the fusion carrying the wild-type *ppc* sequence, none of the point mutations resulted in altered Ppc-GFP expression, as judged by fluorescence from colonies (Fig. 5C). This is in agreement with Ppc being essential for growth on M9-glu; inactivating mutations would not have sustained growth in the suppressor screen. By contrast, OmrA overexpression resulted in strongly reduced fluorescence from the wild-type *ppc-gfp* fusion, whereas each of the point mutant variants was insensitive to inhibition. Quantification of GFP fluorescence from liquid cultures showed a 13-fold decrease in *ppc* translation upon OmrA overexpression (Fig. 5D), but much less in the mutant *ppc-gfp* fusions. Overexpression of OmrB also decreased *ppc-gfp* translation, albeit to a smaller extent.

**Figure 5.**
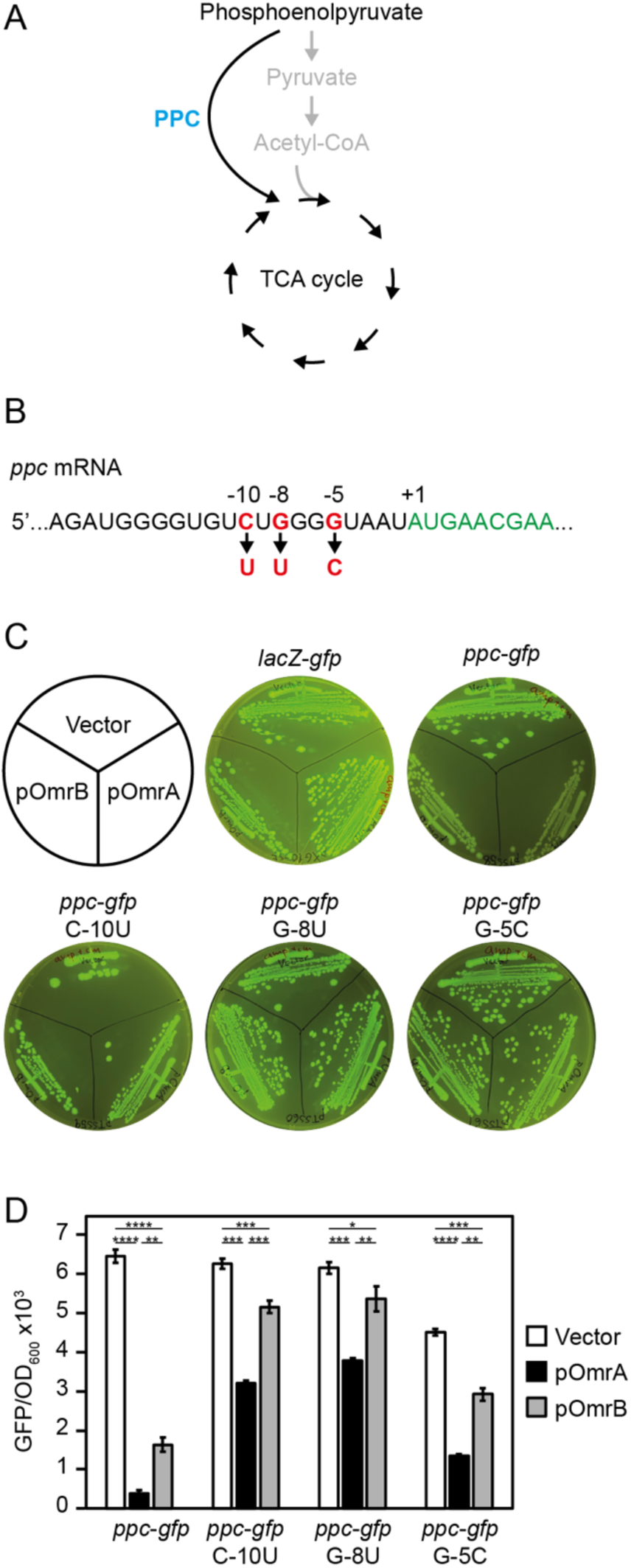
OmrA inhibits synthesis of the metabolic enzyme Ppc. (A) Ppc replenishes the TCA cycle with oxaloacetate by direct conversion from phosphoenolpyruvate. (B) Three independent mutations that suppress OmrA-dependent growth deficiency were identified in the 5’UTR of the *ppc* mRNA. Numbering is relative to the first nucleotide in the *ppc* ORF. (C) Plasmids expressing a translational *ppc-gfp* fusion, or the indicated mutants thereof, were transformed into MG1655 Δ*omrAB* harboring plasmids overexpressing OmrA, OmrB, or an empty control plasmid, respectively, and streaked on Luria agar plates. The plates were placed on a blue-light table to visualize GFP fluorescence and photographed. A translational *lacZ-gfp* fusion plasmid was used as a positive control for GFP fluorescence, and a negative control for OmrA or OmrB regulation. (D) The strains used in C were grown in LB medium in a 96-well plate reader for 16 hours, during which GFP fluorescence and OD_600_ was monitored. Bars show mean values of end-point measurements from three biological replicates. Error bars show standard deviations. Statistical significance was determined using a two-tailed *t* test (\*\*\*\**P* < 0.0001; \*\*\**P* < 0.001; \*\**P* < 0.01; \**P* < 0.05).

### The ppc mRNA is a direct target of OmrA and OmrB

The first six nucleotides in the 5’ tail of OmrA and OmrB are complementary to the segment of the *ppc* 5’UTR harboring the suppressor mutations, suggesting that OmrA/B-dependent repression of *ppc-gfp* expression relies on a base-pairing interaction. To test this, a point mutation (M1:C3G) was introduced in both sRNAs (Fig. 6A). In line with the *ppc-gfp* fusion results, endogenous Ppc-FLAG levels were decreased upon overexpression of OmrA or OmrB, with OmrA giving the strongest effect (Fig. 6B). By contrast, mutant sRNAs harboring the M1 mutation were unable to repress Ppc expression. In line with this, mutation M1, as well as M2 (C1G+C2G), completely abolished OmrA-dependent repression of *ppc-gfp* expression (Fig. 6C). In addition to the predicted base-pairing between the 5’ end of OmrA and *ppc,* a second duplex region could potentially be obtained through a distal interaction involving nts 43-52 (Fig. 6A). Due to its different middle region compared to OmrA, OmrB cannot confer this extended base-pairing (Supplementary Fig. S1). To test whether the downstream region of OmrA contributes to regulation of *ppc*, nts 43-45 of OmrA were mutated to disrupt the predicted base-pairing (M3, Fig. 6A). Similar to the M1 and M2 mutations, the M3 mutation completely abolished regulation of the *ppc-gfp* fusion (Fig. 6C), suggesting that both positions 1-6 and 43-52 may form duplexes with *ppc.* To further test the predicted interaction, we designed a compensatory mutation in *ppc* (M1*: G-8C) which should fully restore complementarity to OmrA-M1 (Fig. 6A). Assaying wild-type and mutant OmrA-*ppc* combinations showed that only cognate pairs (WT-WT or M1-M1*) retained OmrA-dependent repression of *ppc-gfp*, while non-cognate combinations did not (Fig. 6D). To further assess the direct interaction between *ppc* and the sRNAs, we performed gel shift binding assays with *in vitro* transcribed and radioactively labelled *ppc* mRNA and unlabelled OmrA or OmrB, respectively, in the presence of purified Hfq. The presence of OmrA or OmrB resulted in a concentration-dependent slower migration of *ppc* mRNA, indicative of sRNA-mRNA complex formation. Consistent with OmrA showing stronger growth inhibition (Fig. 1B) and Ppc repression (Figs. 5C-D and 6B) than OmrB, the binding affinity for OmrA-*ppc* was higher than for OmrB-*ppc* (Fig. 6E). The location of the OmrA binding site just upstream of the *ppc* start codon suggested the mechanism of regulation to involve inhibition of translation initiation. This was tested in a toe-print assay. A reverse-transcription stop indicative of 30S binding was observed 14 nt downstream of the *ppc* start codon (Fig. 6F). OmrA alone did not affect the toe-print signal. However, in the presence of Hfq, addition of OmrA resulted in a partially inhibited toe-print. This was specific to OmrA, as the unrelated sRNA ArcZ did not affect the toe-print signal, irrespective of the presence of Hfq (Fig. 6F). Together, these results strongly indicate that OmrA and, to a lesser extent, OmrB bind to a complementary sequence in the *ppc* 5’UTR, in a Hfq-dependent manner, to inhibit translation initiation.

**Figure 6.**
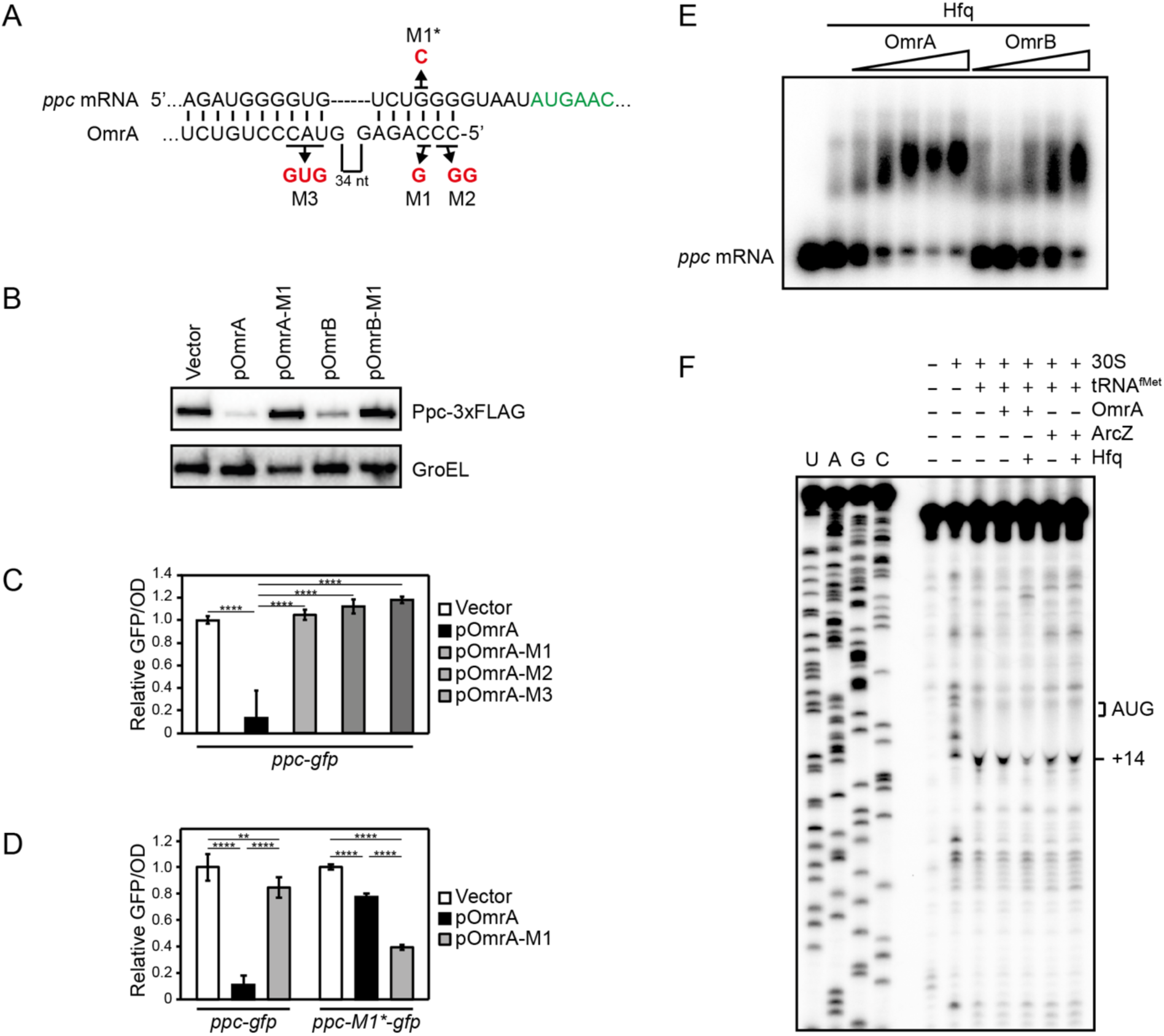
OmrA inhibits Ppc synthesis by direct binding to the *ppc* mRNA. (A) Predicted base-pairing interaction between OmrA and *ppc* 5’UTR. Introduced mutations are indicated and highlighted in red. (B) Western blot analysis of endogenous Ppc-3xFLAG levels in MG1655 Δ*omrAB* upon overexpression of OmrA, OmrB, or the indicated mutants RNAs. Detection of the GroEL protein was used as loading control. (C) GFP fluorescence from MG1655 Δ*omrAB* harboring a translational *ppc-gfp* fusion and an OmrA overexpression plasmid, or the indicated mutant variants, after sixteen hours of growth in a plate reader. (D) GFP fluorescence from MG1655 Δ*omrAB* harboring a translational *ppc-gfp* fusion, or the indicated mutant versions, and an OmrA overexpression plasmid, or mutant variants, after sixteen hours of growth in a plate reader. Bars in C and D: mean values from three biological replicates. Error bars show standard deviations. Statistical significance was determined using a two-tailed *t* test (\*\*\*\**P* < 0.0001; \*\*\**P* < 0.001; \*\**P* < 0.01; \**P* < 0.05). (E) Electromobility assay of 5’ radioactively labelled *ppc* mRNA in the absence or presence of Hfq, OmrA, and OmrB. (F) Toe-print analysis to monitor 30S pre-initiation complex formation on the *ppc* mRNA in the absence or presence of OmrA and/ or Hfq. The sRNA ArcZ served as a non-specific control. A reverse transcription stop at position +14 downstream of the *ppc* start codon upon pre-initiation complex formation is indicated.

## Discussion

The results presented here demonstrated that the OmrA-dependent growth defect in glucose minimal medium could be restored by addition of certain amino acids, pinpointing effects of the sRNA on genes connected to the tricarboxylic acid (TCA) cycle (Fig. 2). In an unbiased mutational screen, chromosomal mutations were obtained that abolished OmrA toxicity. Most mutations mapped to *hfq*, encoding the RNA chaperone that is required for regulation by most enterobacterial sRNAs (Fig. 4B, Supplementary Table S4), suggesting that OmrA-dependent toxicity relies on antisense-based regulation of one or several mRNA targets. The screen identified mutations in several additional genes. The mechanistic basis for a putative phenotype-suppressing effect of most of these mutations is so-far not clear. Strikingly, however, three additional independent mutations were clustered within the ribosome binding site of *ppc* (Fig. 5B), a region of recognizable complementarity to OmrA (Fig. 6A). This gene encodes Ppc, an enzyme that replenishes oxaloacetate in the TCA cycle (Coomes *et al*., 1985). Congruent with the Hfq requirement and the position of the *ppc* mutations, we propose that toxicity is caused by OmrA inhibiting translation of *ppc* mRNA. This mechanism is strongly supported by RNA gel-shifts, translational fusion experiments, and ribosome toeprint assays, using wild-type or mutant OmrA/ B variants as controls (Figs. 5B and C, Fig. 6). Notably, OmrA is far more effective than OmrB in all *in vivo* and *in vitro* experiments, likely because the different middle region of the latter does not promote efficient binding (Supplementary Fig. S1). Thus, this case study of RNA toxicity has identified a target whose known activity can fully account for the physiological effect that results in growth inhibition.

What is the potential impact of these findings regarding physiology? Firstly, as in many other studies, we used sRNA overexpression to observe clear phenotypes. Under the conditions employed, OmrA inhibits *ppc* mRNA, as shown in the experiments reported (Fig. 5), and identification of *ppc* as a direct target is congruent with all growth effects seen in Figures 1-3. Whether this is physiologically relevant is however not yet clear. Subsequent studies are needed to assess more specifically under which conditions chromosomally encoded OmrA may become sufficiently upregulated to account for the regulation observed. Additional strategies could involve competition experiments between wild-type, Δ*omrA*, Δ*omrB*, Δ*omrAB* strains in different media, since they are sufficiently sensitive to reveal even small growth effects, whether they are steady-state- or kinetics-driven. E.g., Spot 42 is one leg of a coherent feed forward loop together with the transcription factor CRP that, in the absence of glucose, binds cAMP and upregulates genes needed for growth on non-preferred carbon sources. The involvement of this sRNA alters the dynamics of regulation, even in single-copy condition (Beisel and Storz, 2011). The involvement of OmrA and B in motility, biofilm, and outer membrane protein synthesis is already well-established (Guillier and Gottesman, 2006; Guillier and Gottesman, 2008; Holmqvist *et al*., 2010; Holmqvist *et al*., 2013; Brosse *et al*., 2016; Jagodnik *et al*., 2017; Hoekzema *et al*., 2019; Romilly *et al*., 2020). If OmrA were to have a role in energy metabolism, this would add to the growing list of sRNAs that affect the flux through metabolic pathways (e.g. SgrS, Spot 42, SdhX, and others (Papenfort and Storz, 2024)).

A second issue concerns what RNA toxicity implies, and which kind of RNAs are likely to confer growth effects under certain conditions – not necessarily during artificial overexpression. It is known that mutant versions of RNAs can interfere with vital functions (Sicot and Gomes-Pereira, 2013; Mittal *et al*., 2018). One can also imagine that even normal chromosomally encoded RNAs may have such effects, and sRNAs in particular are good candidates. A sufficient degree of complementarity to a certain mRNA may often only result in insignificant effects, undetectable as toxicity. At higher levels, however, as generally obtained in overexpression experiments, biologically meaningful targets may be revealed. More trivial explanations of RNA toxicity may involve off-target effects. I.e., very high sRNA levels may partially overcome poor complementarity with targets whose gene products are essential. In principle, any RNA can therefore be toxic under some set of conditions. So far, RNAs that are toxic due to sequestration of essential proteins have, to our knowledge, so far not be found. By contrast, antisense-type sRNAs have properties that match requirements for condition-dependent toxic effects. Most sRNAs are regulatory hubs with large target repertoires (Beisel and Storz, 2010), and perturbations may therefore be harmful under some conditions. Thus, what can be considered cases of misregulation may help to identify relevant roles of a “toxic” RNA under different conditions. Whether this applies for OmrA/ *ppc* needs to be established.

A starting point for this study was our interest in OmrA/B as sibling RNAs, asking the question whether these paralogous RNAs exclusively regulate the same set of target mRNAs, or whether their evolutionary trajectory causes them to “speciate” in terms of separate targeting. So far, only one study has demonstrated this; OmrA rather than OmrB represses translation of the *btuB* mRNA (Bastet *et al*., 2024). In our experiments, we also see a clear difference in targeting of *ppc* mRNA, again with OmrA being by far more effective. Sibling sRNAs have attracted much interest, and many pairs (sometimes more than two) have been reported (for a review, see (Caswell *et al*., 2014)). Some sRNAs, as OmrA and B, are encoded in tandem. Some have independent genes. Though the list of siblings grows and includes many bacterial species, it is yet difficult to assess clear patterns. It seems that redundancy with respect to targets is quite common. However, in a few cases, siblings have evolved apart and, in some cases, are also under separate transcriptional regulation. E.g., the two AbcR sRNA paralogs in *Agrobacterium tumefaciens* act on different targets; AbcR1 alone regulates multiple ABC transporters (Overlöper *et al*., 2014). Though the implications of sibling sRNAs in bacteria is still elusive, it is likely that they follow a pattern that is well-known for other duplicated genes. Paralogs are free to evolve separate functions, as exemplified by two-component systems (Capra and Laub, 2012). In the OmrA/B case, the significant sequence changes in the middle region of the RNA may suggest that more distinct and separate targets for either of these RNAs will be discovered.

Even though this study clearly identifies *ppc* mRNA as a direct OmrA target under overexpression conditions, it remains to be established whether comparable regulation occurs at endogenous OmrA levels. Addressing this will require determining how OmrA abundance varies across growth states and assessing the consequences of *ppc* repression in single-copy genomic contexts. Importantly, Ppc activity depends not only on protein concentration but also on its tetrameric assembly and extensive allosteric regulation (Corwin and Fanning, 1968; Kai *et al*., 1999). Thus, even modest OmrA-dependent reductions in *ppc* translation may have amplified effects on TCA-cycle flux. More broadly, these findings add to the growing appreciation that bacterial sRNAs can directly influence central metabolism (Papenfort and Storz, 2024). Moreover, OmrA and OmrB may offer an informative example of how sibling sRNAs diverge to mediate distinct regulatory outputs. Finally, the OmrA–*ppc* interaction uncovered here illustrates how “toxic” overexpression phenotypes can reveal biologically relevant sRNA–mRNA connections that would otherwise remain cryptic.

## Acknowledgements

We thank the research group for valuable input and fruitful discussions.

## Supplementary data

Supplementary data is available at NAR online.

