## Supplementary Information for "OmrA sRNA Inhibits Translation of Phosphoenolpyruvate Carboxylase to Impair TCA-Cycle Flux"

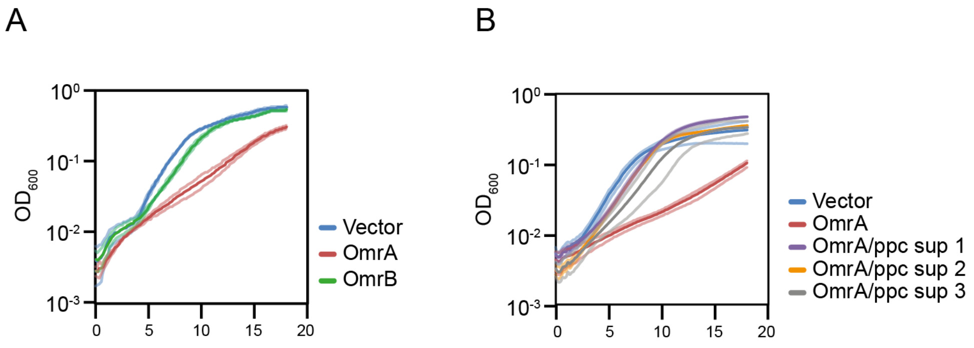


Figure S1. (A) *E. coli* MG1655 Δ*omrAB* harboring an empty vector, pOmrA, or OmrB, was grown in M9-glucose media in a 96 well plate. (B) Growth of suppressor strains harboring mutations in *ppc* upon OmrA overexpression. Bright colored lines: mean growth rate. Faint lines: standard deviation based on four biological replicates.

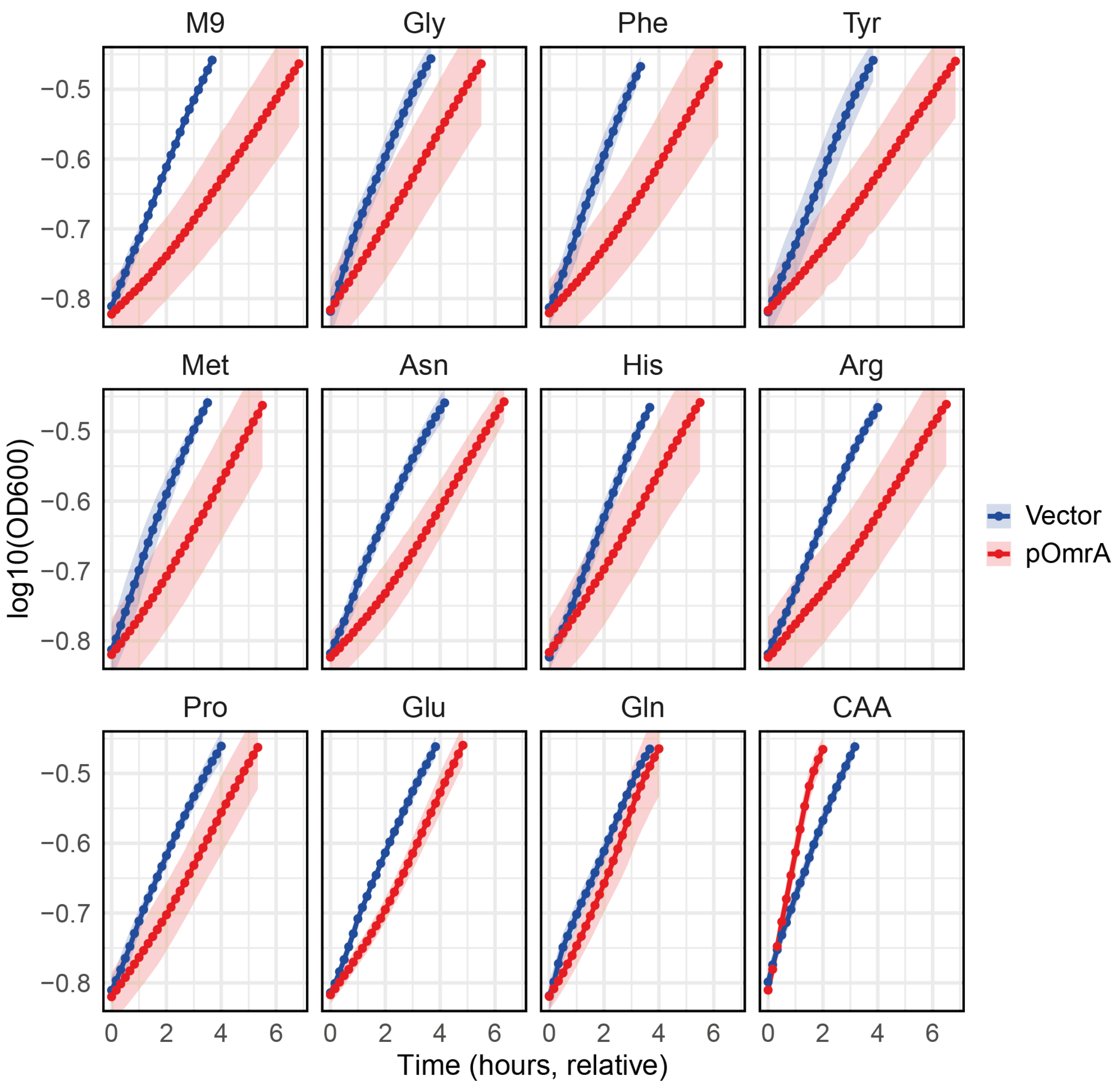


Figure S2. *E. coli* MG1655 Δ*omrAB* harboring an empty vector or pOmrA was grown in M9-glucose media supplemented with the indicated amino acids in a 96 well plate. The exponential growth phase is shown.

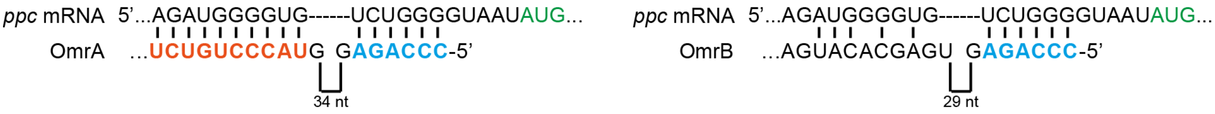


```
OmrA    CCCAGAGGTATTGATTGGTGAGATTATTCGGTACGCTCTTCGTACCCTGTCTCTTGCACCAACCTGCGCGGATGCGCAGGTTTTTTTT
OmrB    CCCAGAGGTATTGATAGGTGAAGTCA------ACTTCGGGTTGAGCACATGAATTACACCAGCCTGCGCAGATGCGCAGGTTTTTTTT
        *************** *****..*.*      ** ..   .  * * ..*   **.*****.*******.*****************
```

Figure S3. Predicted base-pairing between OmrA or OmrB and the *ppc* 5'UTR (top panels). Alignment indicating sequence deviations in the middle regions of the two sRNAs.

Table S1. Oligos used in this study

| Name | Description | Sequence (5'-3') |
|---|---|---|
| EHO-1006 | Fwd for PCR of pZE12-luc blunt (pLlacO-1) | GTTTTTATGCATGTGCCACCTGACGTCTAAGAAAC |
| EHO-1122 | Rev for PCR of pZE12-luc blunt (pLlacO-1) | CGGCGGATTTGTCCTACT |
| EHO-827 | Fwd for colony PCR on pXG10-SF. Used for plasmid ppc-GFP sequencing. | TGGGATATATCAACGGTGGT |
| EHO-828 | Rev for colony PCR on pXG10-SF. Used for plasmid ppc-GFP sequencing. | GAATTGGGACAACTCCAGTG |
| EHO-2287 | Fwd for creating a translational fusion of MG1655 ppc to GFP-SF in pXG10-SF, including the whole 5'UTR and the first 20 codons of the ORF. With NsiI site and 5 nt overhang | GTTTTATGCATGACGACGAAAAGCAAAGCC |
| EHO-2288 | Rev for creating a translational fusion of MG1655 ppc to GFP-SF in pXG10-SF, including the whole 5'UTR and the first 20 codons of the ORF. With NheI site and 5 nt overhang | GTTTTGCTAGCTCCCAGCACTTTGCCGAG |
| EHO-2289 | Fwd for introducing mutation 1 (position -10:C->T) in the ppc-GFP translational fusion plasmid | TGGGGTGTTTGGGGTAATATGAACGAACAATATTCC |
| EHO-2290 | Rev for introducing mutation 1 (position -10:C->T) in the ppc-GFP translational fusion plasmid | TTACCCCAAACACCCCATCTTATCGTTTGATAGCCC |
| EHO-2291 | Fwd for introducing mutation 2 (position -8: G->T) in the ppc-GFP translational fusion plasmid | GGGTGTCTTGGGTAATATGAACGAACAATATTCCGC |
| EHO-2292 | Rev for introducing mutation 2 (position -8: G->T) in the ppc-GFP translational fusion plasmid | TATTACCCAAGACACCCCATCTTATCGTTTGATAGC |
| EHO-2293 | Fwd for introducing mutation 3 (position -5: G->C) in the ppc-GFP translational fusion plasmid | TGTCTGGGCTAATATGAACGAACAATATTCCGCATT |
| EHO-2294 | Rev for introducing mutation 3 (position -5: G->C) in the ppc-GFP translational fusion plasmid | TCATATTAGCCCAGACACCCCATCTTATCGTTTGAT |
| EHO-2307 | Fwd for C-terminal 3xFLAG tagging of ppc, containing region upstream of stop codon of ppc (in capital), and homologous region of 3x-FLAG-FRT-KmR-FRT from pSUB11 (in non-capital). | CACTATTGCCGGGATTGCGGCAGGTATGCGTAATACCGGCgactacaaagaccatgacgg |
| EHO-2308 | Rev for C-terminal 3xFLAG tagging of ppc, containing region upstream of stop codon of ppc (in capital), and homologous region of 3x-FLAG-FRT-KmR-FRT from pSUB11 (in non-capital). | ACCCTCGCGCAAAAGCACGAGGGTTTGCAGAAGAGGAAGAcatatgaatatcctccttag |
| EHO-2309 | Fwd to MG1655 ORF of ppc. Used for PCR screen of 3xFLAG insertion on C-term of ppc | GAAGACATCAAAGTGGTGCTG |
| EHO-2310 | Rev to region downstream of MG1655 ppc ORF. Used for PCR screen of 3xFLAG insertion on C-terminal of ppc | AATGGCCTGTAGCAGAGTAGA |
| EHO-2312 | Fwd for T7 for ppc gene. Used for in vitro transcription. | gaaattaatacgactcactataGGACGACGAAAAGCAAAGCCCGAG |
| EHO-2313 | Rev for T7 for ppc gene. Used for in vitro transcription. | TACTGACATTACTACGCAATGCGGAA |
| EHO-2470 | Rev for toeprint assay of ppc gene | TTCTCCCAACGCATCCTTGATGGTTT |

| | | |
|---|---|---|
| EHO-2479 | Fwd for introducing mutation ppc M1*(position -9: GUC -> CUC) in the ppc-GFP translational fusion plasmid | GTCTCGGGTAATATGAACGAACAATATTC |
| EHO-2480 | Rev for introducing mutation ppc M1*(position -9: GUC -> CUC) in the ppc-GFP translational fusion plasmid | ACCCGAGACACCCCATCTTATC |
| EHO-2234 | Fwd T7-arcZ, anneal to EHO-2235 | gtttttttaatacgactcactataggTTTCCCTGGTGTTGGCGCAGTATTCGCGCACCCCGGTCAAACCGGGGTCATTTTTT |
| EHO-2235 | Rev T7-arcZ, anneal to EHO-2234 | AAAAAATGACCCCGGTTTGACCGGGGTGCGCGAATACTGCGCCAACACCAGGGAAAcctatagtgagtcgtattaaaaaaac |
| EHO-2370 | Fwd for introducing mutation omrA M3 (position 43: TAC -> GTG) into pEH67 | TCGGTGCCTGTCTCTTGC |
| EHO-2371 | Rev for introducing mutation omrA M3 (position 43: TAC -> GTG) into pEH67 | AGGCACCGAAGAGCGTAC |
| EHO-245 | Fwd for FRT-Tet-FRT ampli with omrB upstream homology (in capital), for omrAB deletion with lambda red | CCCGTTGGTTCAAGGGGTGGCGTGTTTTCATCGTGGGAATgaattcgagctcggtacccg |
| EHO-246 | Rev for FRT-Tet-FRT ampli with omrA homology (in capital), for omrAB deletion with lambda red | CCTGCGCATCCGCGCAGGTTGGTGCAAGAGACAGGGTACgctatgaccatgattacgcc |
| EHO-247 | Fwd upstream of omrAB deletion, 5' junction PCR | CTTAGTGGGTAAACGCTTACATTC |
| EHO-249 | Rev downstream of omrAB deletion, 3' junction PCR | CAACTCCCTTTGCTCTGATTCAG |
| EHO-2528 | For T7-Hammerhead-omrA M1, anneal to EHO-2529, fill-in | gaaattaatacgactcactataGGGAGATGGGCTGATGAGTCCGTGAGGACGAAACGGTACCCGGTACCGTCCCGAGAGGTATTGATTGG |
| EHO-2529 | Rev T7-Hammerhead-omrA M1, anneal to EHO-2528, fill-in | AAAAAAAACCTGCGCATCCGCGCAGGTTGGTGCAAGAGACAGGGTACGAAGAGCGTACCGAATAATCTCACCAATCAATACCTCTCGGGA |
| EHO-406 | OmrA northern probe | GAGACAGGGTACGAAGAGCGTACCGAATAATCTCACC |
| EHO-407 | OmrB northern probe | GTGTAATTCATGTGCTCAACCCGAAGTTGACTTCACC |
| EHO-867 | tmRNA northern probe | TGGTGGAGCTGGCGGGAGTT |

Table S2. Bacterial strains used in this study

| **Strain number** | **Name** | **Genotype** | **Plasmid** | **Reference** |
|---|---|---|---|---|
| | E. coli TOP10 | F- mcrA Δ( mrr-hsdRMS-mcrBC) Φ80lacZΔM15 Δ lacX74 recA1 araD139 Δ( araleu)7697 galU galK rpsL (StrR) endA1 nupG | | |
| EHS-4092 | E. coli TOP10 | F- mcrA Δ( mrr-hsdRMS-mcrBC) Φ80lacZΔM15 Δ lacX74 recA1 araD139 Δ( araleu)7697 galU galK rpsL (StrR) endA1 nupG | pTSS58 | this study |
| EHS-4102 | E. coli TOP10 | F- mcrA Δ( mrr-hsdRMS-mcrBC) Φ80lacZΔM15 Δ lacX74 recA1 araD139 Δ( araleu)7697 galU galK rpsL (StrR) endA1 nupG | pTSS59 | this study |
| EHS-4103 | E. coli TOP10 | F- mcrA Δ( mrr-hsdRMS-mcrBC) Φ80lacZΔM15 Δ lacX74 recA1 araD139 Δ( araleu)7697 galU galK rpsL (StrR) endA1 nupG | pTSS60 | this study |
| EHS-4104 | E. coli TOP10 | F- mcrA Δ( mrr-hsdRMS-mcrBC) Φ80lacZΔM15 Δ lacX74 recA1 araD139 Δ( araleu)7697 galU galK rpsL (StrR) endA1 nupG | pTSS61 | this study |

| | | | | |
|---|---|---|---|---|
| EHS-4145 | E. coli TOP10 | F- mcrA Δ( mrr-hsdRMS-mcrBC) Φ80lacZΔM15 Δ lacX74 recA1 araD139 Δ( araleu)7697 galU galK rpsL (StrR) endA1 nupG | pKB009 | this study |
| KBS-0100 | E. coli TOP10 | F- mcrA Δ( mrr-hsdRMS-mcrBC) Φ80lacZΔM15 Δ lacX74 recA1 araD139 Δ( araleu)7697 galU galK rpsL (StrR) endA1 nupG | pKB013 | this study |
| EHS-3164 | MG1655 | Δ *omrAB*::FRT-TetR-FRT | pJV300 | this study |
| EHS-3165 | MG1655 | Δ *omrAB*::FRT-TetR-FRT | pEH67 | this study |
| EHS-3166 | MG1655 | Δ *omrAB*::FRT-TetR-FRT | pEH68 | this study |
| EHS-3833 | MG1655 | Δ *omrAB*::FRT-FRT | pJV300 | this study |
| EHS-3834 | MG1655 | Δ *omrAB*::FRT-FRT | pEH67 | this study |
| EHS-3835 | MG1655 | Δ *omrAB*::FRT-FRT | pEH68 | this study |
| EHS-4020 | MC4100 relA- | | pJV300 | (1) |
| EHS-4021 | MC4100 relA- | *hfq-F42A* | pJV300 | (1) |
| EHS-4022 | MC4100 relA- | *hfq-Y25D* | pJV300 | (1) |
| EHS-4023 | MC4100 relA- | *hfq-R16A* | pJV300 | (1) |
| EHS-4026 | MC4100 relA- | | pEH67 | (1) |
| EHS-4027 | MC4100 relA- | *hfq-F42A* | pEH67 | (1) |
| EHS-4028 | MC4100 relA- | *hfq-Y25D* | pEH67 | (1) |
| EHS-4029 | MC4100 relA- | *hfq-R16A* | pEH67 | (1) |
| EHS-4093 | MG1655 | Δ *omrAB*::FRT-FRT | pXG-0, pJV300 | this study |
| EHS-4095 | MG1655 | Δ *omrAB*::FRT-FRT | pTSS58, pJV300 | this study |
| EHS-4098 | MG1655 | Δ *omrAB*::FRT-FRT | pTSS58, pEH67 | this study |
| KBS-0081 | MG1655 | Δ *omrAB*::FRT-FRT | pTSS58, pEH171 | this study |
| KBS-0082 | MG1655 | Δ *omrAB*::FRT-FRT | pTSS58, pEH80 | this study |
| KBS-0085 | MG1655 | Δ *omrAB*::FRT-FRT | pTSS58, pKB009 | this study |
| KBS-0106 | MG1655 | Δ *omrAB*::FRT-FRT | pKB013, pJV300 | this study |
| KBS-0107 | MG1655 | Δ *omrAB*::FRT-FRT | pKB013, pEH67 | this study |
| KBS-0108 | MG1655 | Δ *omrAB*::FRT-FRT | pKB013, pEH171 | this study |
| EHS-4242 | MG1655 | *ppc*-3xFLAG (C-terminal) | | this study |
| EHS-4243 | MG1655 | Δ *omrAB*::FRT-FRT *ppc*-3xFLAG (C-terminal) | | this study |
| KBS-0165 | MG1655 | Δ *omrAB*::FRT-FRT *ppc*-3xFLAG (C-terminal) | pJV300 | this study |
| KBS-0166 | MG1655 | Δ *omrAB*::FRT-FRT *ppc*-3xFLAG (C-terminal) | pEH67 | this study |
| KBS-0167 | MG1655 | Δ *omrAB*::FRT-FRT *ppc*-3xFLAG (C-terminal) | pEH171 | this study |
| KBS-0168 | MG1655 | Δ *omrAB*::FRT-FRT *ppc*-3xFLAG (C-terminal) | pEH68 | this study |
| KBS-0169 | MG1655 | Δ *omrAB*::FRT-FRT *ppc*-3xFLAG (C-terminal) | pEH172 | this study |

Table S3. Plasmids used in this study

| Name | Description | Marker | Reference |
|---|---|---|---|
| pJV300 | pZE12-luc based control plasmid, -1 site religated to second position of XbaI site, destroys XbaI site (colE1, Amp), PLlacO promoter reads into rrnB terminator | Amp | (2) |
| pXG-0 | Control plasmid for cellular autofluorescence | Cm | (3) |
| pXG10-SF | pXG-10 derivative with sfgfp, for translational sfGFP fusions | Cm | (4) |
| pSIM5-Tet | λ red expression plasmid | Tet | (5) |
| pTSS58 | *ppc-gfp* translational fusion. The leader of ppc (from TSS until 20 codons into the gene) was cloned inframe with the GFP in pXG10-SF using NsiI and NheI RE sites. | Cm | this study |
| pTSS59 | *ppc M1-gfp* translational fusion. The leader of ppc with a C->T mutation at the -10 position (from TSS until 20 codons into the gene) was cloned inframe with the GFP in pXG10-SF using NsiI and NheI RE sites. | Cm | this study |
| pTSS60 | *ppc M2-gfp* translational fusion. The leader of ppc with a G->T mutation at the -8 position (from TSS until 20 codons into the gene) was cloned inframe with the GFP in pXG10-SF using NsiI and NheI RE sites. | Cm | this study |
| pTSS61 | *ppc M3-gfp* translational fusion. The leader of ppc with a G->C mutation at the -5 position (from TSS until 20 codons into the gene) was cloned inframe with the GFP in pXG10-SF using NsiI and NheI RE sites. | Cm | this study |
| pKB013 | based on pTSS58, with *ppc M1** compensatory (EHO-2479/2480) to OmrA M1 | Cm | this study |
| pEH67 | OmrA cloned in pZE12-luc blunt (pLlacO-1) | Amp | (6) |
| pEH68 | OmrB cloned in pZE12-luc blunt (pLlacO-1)/XbaI | Amp | (6) |
| pEH171 | based on pEH67, with OmrA M1 (EHO-2370/2371) mutated position 3-C -> G | Amp | (7) |
| pEH80 | based on pEH67, with OmrA M2 mutated position1-2: CC -> GG | Amp | (7) |
| pKB009 | based on pEH67, with OmrA M3 (EHO-2370/2371) mutated position 43-TAC -> GTG | Amp | this study |
